# Development of an MRI Compatible Nasal Drug Delivery Method: Probing Nicotine Addiction Dynamics

**DOI:** 10.1101/2020.09.17.302703

**Authors:** Lilianne R. Mujica-Parodi, Rajat Kumar, Michael Wenke, Anar Amgalan, Andrew Lithen, Sindhuja T. Govindarajan, Rany Makaryus, Helene Benveniste, Helmut H. Strey

## Abstract

**Background:** Substance abuse is a fundamentally dynamic disease, characterized by repeated oscillation between craving, drug self-administration, reward, and satiety. To model nicotine addiction as a control system, an MR-compatible nicotine delivery system is needed to elicit cyclical cravings.

**Method:** Using a concentric nebulizer, inserted into one nostril, we delivered each dose—each equivalent to a single cigarette puff—using a syringe pump by nebulizing the nicotine solution using pressurized medical air. A control mechanism permits dual modes: one delivers puffs on a fixed interval programmed by researchers; with the other, subjects press a button to self-administer each nicotine dose. Subjects were therefore able to intuitively “smoke” the equivalent of a cigarette, one “puff” at a time. We dosed each “puff” such that one cigarette would be equal, in nicotine content, to 10 puffs.

**Results:** We tested the viability of this delivery method for studying the brain’s response to nicotine addiction in three steps. First, we established the pharmacokinetics of nicotine delivery, using a dosing scheme designed to gradually achieve saturation, as with a cigarette. Second, we lengthened the time between micro-doses to elicit craving cycles, using both fixed-interval and subject-driven behavior. Finally, we confirmed that the fixed-interval protocol reliably activates brain circuits linked to addiction.

**Conclusion:** Our MR-compatible nasal delivery method enables the measurement of neural circuit responses to drug doses on a single-subject level, allowing the development of data-driven predictive models to quantify individual dysregulations of the reward control circuit causing addiction.

## Introduction

Nicotine is the most common drug of abuse in the United States(Medicine 2010), with addiction strength comparable to cocaine, heroin, and alcohol(“How Tobacco Smoke Causes Disease: The Biology and Behavioral Basis for Smoking-Attributable Disease: A Report of the Surgeon General” 2010; Abuse 2009). It is the primary addictive component of tobacco, and its use markedly increases risk for cancer, heart disease, asthma, miscarriage, and infant mortality(“How Tobacco Smoke Causes Disease: The Biology and Behavioral Basis for Smoking-Attributable Disease: A Report of the Surgeon General” 2010). Excitatory vs. inhibitory control plays a key role in addictive behavior; for example, decreasing glutamate or increasing GABA transmission blunts nicotine craving in rats (D’Souza and Markou 2013; Markou 2008). Importantly, however, substance abuse is a fundamentally *dynamic* disease, characterized by repeated oscillation between craving, drug self-administration, reward, and satiety. While there is a general appreciation for the heuristic value of separating out these stages(Koob and Volkow 2016), current clinical research has primarily focused on identifying nodes and causal connections within the meso-circuit of interest, but has yet to take the next step in treating these nodes and connections as a self-interacting dynamical system evolving over time(Mujica-Parodi, Cha, and Gao 2017). The value of a dynamical systems approach is the potential for predicting trajectories for addiction as well as recovery. These trajectories are likely to be nonlinear (e.g., involving thresholds, saturation, and self-reinforcement) and individual-specific.

As a first step towards this approach, we identified several requirements for nicotine delivery. First, the delivery method should be able to mimic the pharmacokinetics of the addictive substance, as there is significant evidence that absorption speed affects cravings (Le Houezec 2003). Second, delivery should be capable of eliciting multiple cycles that transition between craving, resistance, breakdown of resistance, and satiety, as a prerequisite to data-driven computational modeling of feedback within the control circuit. Third, craving cycles—and associated behavioral self-administration—should be triggered solely on drug pharmacokinetics and their dynamic interactions with the brain. This has the advantage of decoupling chemical effects from cigarette-specific motor, olfactory, and visual cues, each of which may trigger its own responses in the brain. Fourth, to permit investigation of the relationship between drug-seeking behavior and its underlying neurobiology, it should allow subjects to self-administer nicotine. Fifth, the device should be MR-compatible, as fMRI is currently the only non-invasive neuroimaging tool with subcortical-cortical coverage of the entire prefrontal-limbic-reward circuit. While neuroimaging has historically relied on heavily fitted and filtered amplitude and correlational statistics, recent technological developments in ultra-high-field (7T) fMRI and simultaneous multi-slice pulse sequences, provide sufficient signal/noise to permit single subject, non-trial-averaged, and sub-second time-resolution time-courses that retain much greater dynamic information(DeDora et al. 2016).

To administer nicotine within a controlled environment, researchers have employed one of five strategies. These include: (1) *smoking* (either directly, or through an apparatus that ensures controlled delivery), (2) *intravenous* (either as a bolus injection or continuously by drip), (3) *transdermal* (using a nicotine patch), (4) *absorption through the lungs* (nicotine solution nebulized in mouth), or (5) *absorption through the mucus membranes* (nicotine solution sprayed in nose/throat) (Stead et al. 2012).

Several of those strategies introduce additional practical problems and scientific confounds. Although other groups have designed MR-compatible smoke delivery devices for use in 3T scanners (Frederick et al. 2007), this method can raise environmental safety concerns regarding second-hand smoke inhalation and lingering odor. Moreover, tobacco is a natural substance with non-standardized composition, and thus prevents precise control of dosage. Intravenous bolus is invasive with inherent greater risks for overdose/toxicity and has pharmacokinetics different from those of smoking. Indeed, in order to satisfy cravings nicotine injections need to achieve significantly higher doses than cigarette puffs(Rose et al. 1999), which may be related to the nicotine bolus being diluted by having to pass through the heart and liver. Nicotine patches are noninvasive and minimize discomfort from nicotine. However, their extended release of nicotine also renders their pharmacokinetics different from smoking, and do not elicit cravings. Inhaling nebulized nicotine by mouth would intuitively seem to approximate the act of smoking, but again the pharmacokinetics do not translate well to cigarettes. The delivery efficiency of nebulizers depends strongly on how deeply liquid droplets can penetrate the lung, which itself is determined by size and aerosol flow rate (Phipps, Pharm, and Gonda 1990). Delivery by e-cigarettes uses nicotine concentrations in their solutions that are much higher than those of cigarettes because of the relative inefficiency of the lung to absorb liquid droplets. Heating can miniaturize the droplets; however, heating elements are electric, and therefore are not MR-compatible.

Nasal delivery of nicotine addresses many of the problems described above. Nasal sprays produce pharmacokinetics much closer to that of cigarettes, as mucus membranes in the nasal cavities deliver drugs more directly into the brain (Crowe et al. 2018). This may explain why the commercial nicotine replacement medication Nicotrol NS (Pfizer), delivered through the nose, shows significantly greater nicotine craving relief than the same Nicotrol inhaler inhaled through the mouth (Stead et al. 2012). Moreover, because nasal sprays do not trigger motor, visual, or olfactory cues associated with smoking, this mode of delivery permits researchers to more precisely isolate the addiction dynamics of nicotine alone.

To exploit the advantages of nasal nicotine delivery, we designed a delivery device to operate as a concentric nebulizer, inserted into one nostril. We delivered each dose—each equivalent to a single cigarette puff—using a syringe pump and nebulized the dose using pressurized medical air. A control mechanism permits dual modes: one delivers puffs on a fixed interval programmed by researchers; with the other, subjects press a button to self-administer each nicotine dose. Subjects were therefore able to intuitively “smoke” the equivalent of a cigarette, one “puff” at a time. We dosed each “puff” such that one cigarette would be equal, in nicotine content, to 10 puffs. We then tested the viability of this delivery method for studying the brain’s response to nicotine addiction in three steps. First, we established the pharmacokinetics of nicotine delivery, using a dosing scheme designed to gradually achieve saturation, as with a cigarette. Second, we lengthened time between micro-doses to elicit craving cycles, using both fixed-interval and subject-driven behavior. Finally, we confirmed that the fixed-interval protocol reliably activates brain circuits linked to addiction(Koob and Volkow 2016).

## Materials and Methods

### Delivery Apparatus Design

The nicotine delivery apparatus uses a glass concentric mass spectroscopy nebulizer (Meinhart Model TR-30-A1) to deliver Nicotrol nasal spray (NS) into the subject’s nasal cavity. This setup can be used to deliver any water-soluble drug intranasally in an MRI environment **(Figure 1)**. A wide nostril guard (Aptar Pharma) and drawstring strap were used to orient the nebulizer straight in the nose and to ensure the nebulizer does not penetrate too deeply. The nebulizer’s inner capillary contained the Nicotrol NS, which was driven out of the capillary as a mist when medical air is driven through the outer chamber. Medical air pressure was held constant at 15PSI with a regulator (VWR Breathair). This was found to be the optimal air pressure for producing a fine mist of Nicotrol NS that deposits deep enough in the turbinate for rapid absorption while maximizing subject comfort (Cheng et al. 2001). The medical air flowed through a length of Tygon Non-DEHP Food and Beverage Tubing (S3-B-44-3, 3/16”x5/16”, Cole Parmer) and then through an air valve (Clippard MME-32QES, Airoyal NY), which only allowed air to reach the nebulizer during puff delivery. A second air valve was used to quickly depressurize the tubing at the termination of each puff. The Nicotrol NS was dispensed in 10µL puffs, each lasting three seconds, using an automated syringe pump (KDS100, KD Scientific). This syringe pump compressed a Hamilton glass syringe (Fischer Scientific), which was connected to a length of capillary tubing (Meinhard F2-15, 0.8mm inner diameter) and a three-way valve (PEEK 3-Port Flow, V100T, Idex Health Science) before feeding into the nebulizer. Nicotrol NS sat in the length of tubing adjacent to the nebulizer, while the remainder of the tubing feeding back to the syringe pump was filled with cosmetic jojoba oil. This oil was used because it is incompressible and immiscible with the Nicotrol NS, allowing for accurate delivery of a specific volume of Nicotrol.

**Figure 1:**
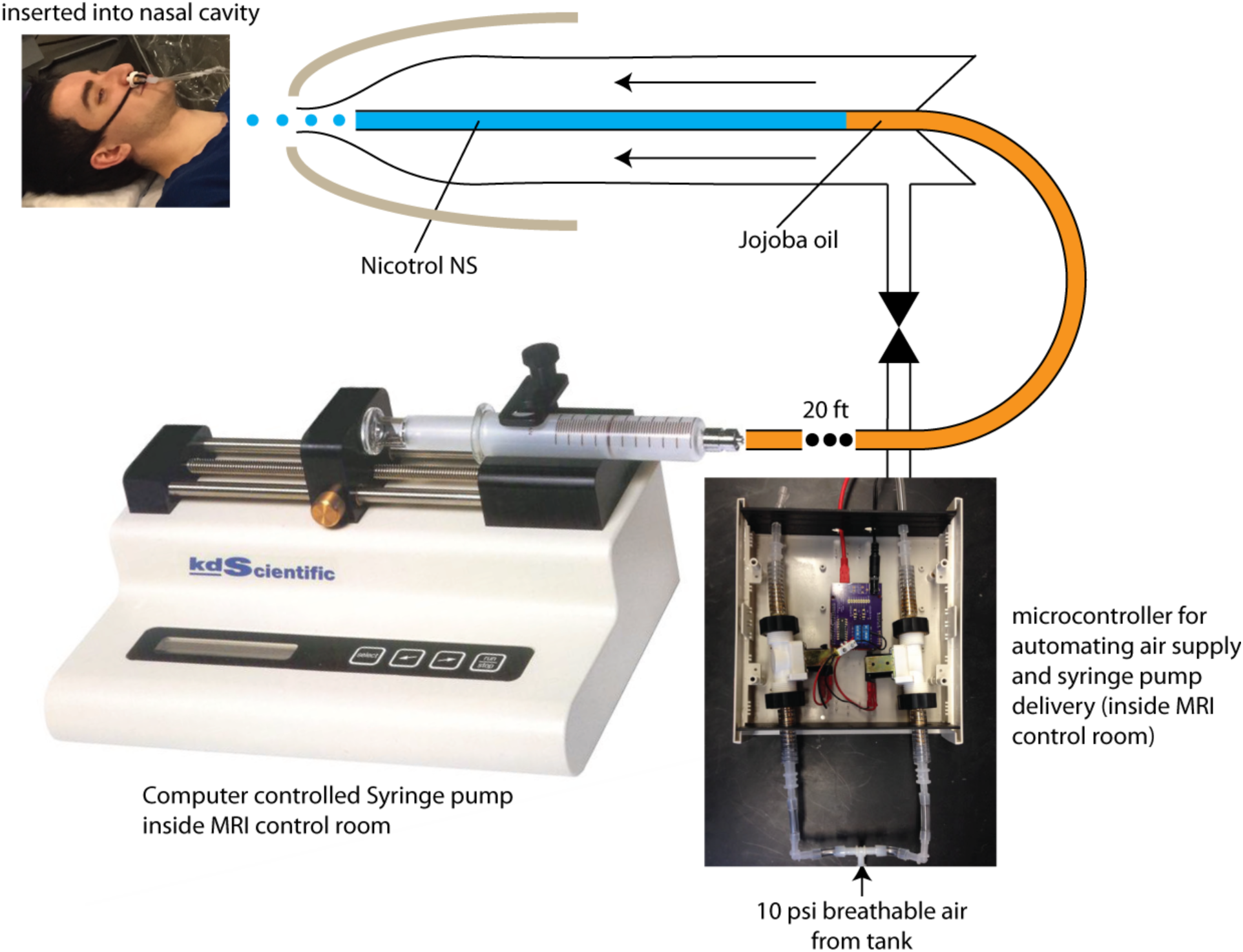
Schematic of the MR-compatible intranasal nicotine delivery apparatus.

An Arduino Uno microcontroller coordinated the release of puffs. The experimental task, which was programmed in MATLAB (R2018a; Mathworks, Natick MA), sent a signal to this microcontroller every time a puff was to be delivered. At this point, the Arduino unit triggered the following events. First, the microcontroller opened the air valve for 16 seconds, allowing air pressure to build up in the nebulizer. A countdown was shown to the subject indicating when nicotine would be delivered. At the end of the countdown, the microcontroller then signaled the syringe pump to compress 10µL over 3 seconds, pushing 10µL of Nicotrol NS out of the nebulizer as a mist. The air valve remained open for an additional 16 seconds after the end of the previous countdown to ensure all Nicotrol at the nebulizer tip was delivered. Finally, the main air valve was closed, and the exhaust valve opened to quickly depressurize the tubes carrying air. The task screen reverted to the cravings indicator.

### Preparing the Apparatus for Experimentation

Immediately prior to use on any subject, clean and sterilized capillary tubing was filled with jojoba oil and Nicotrol while carefully avoiding to trap air in the path to ensure the accuracy of Nicotrol NS delivery. The length of capillary tubing leading to the Hamilton syringe was filled with jojoba oil by submerging the capillary end connecting to the three-way valve in oil; we then used the Hamilton syringe to draw oil to fill the capillary tube. Leaving an oil bead at the end of the capillary tube proximal to the syringe pump, the Hamilton syringe was disconnected, fully compressed, reconnected, and pulled back to fill with oil while excluding any air. A volume of 300µL Nicotrol NS was drawn in an Eppendorf tube and spun down using a micro-centrifuge to remove any air bubbles. This volume contained 3mg nicotine, the amount of nicotine present in three typical cigarettes (the potential for nicotine overdose was thus eliminated since this was the maximum dose that could be delivered). A dyed jojoba oil solution was prepared by mixing 1mL of oil with a few drops of food coloring (Wilton Candy Colors) and homogenizing with a vortex mixer or sonication. The short capillary end of the three-way valve was submerged in the Nicotrol NS. A 1mL draw syringe was connected to the long capillary end of the three-way valve and pulled back until all the Nicotrol NS was drawn into the capillary tube. Leaving a Nicotrol bead at the short capillary end, this end was submerged in the dyed jojoba. The draw syringe was slowly drawn up until the long capillary end was filled. Slow filling was needed to reduce viscous fingering at the interface of the Nicotrol NS and dyed oil. The syringe pump was compressed to produce a bead at the end of the connected capillary tube. This capillary was then screwed into the three-way valve with the valve to the syringe pump closed off. The three-way valve was rotated 90 degrees to close off the capillary leading to the nebulizer and connecting the syringe pump capillary to the short capillary end. This releases the built-up pressure from screwing in the syringe pump capillary. The syringe pump was then compressed until undyed oil was visible in the short capillary tube, indicating no air is present in the three-way valve. The draw syringe was replaced with the nebulizer. The three-way valve was rotated 180 degrees to close off the short capillary end and make a continuous path from the syringe pump to the nebulizer. The syringe pump was then compressed until Nicotrol NS filled the inner capillary of the nebulizer. The nebulizer, nose cone, and strap were sanitized and assembled. The nebulizer was then positioned in the subject’s nostril. At the termination of each experiment, the three-way valve, nebulizer, associated capillary tubes, and nose cone were disinfected by passing through with 20ml soap water, and then 10mL Sporgon disinfectant (Decon Laboratories). The assembly was then submerged in a Sporgon bath for 3 hours, and then rinsed with 20mL deionized water. The assembly tubing was then dried using pressurized medical air.

### Subjects

To establish pharmacokinetics of our delivery mechanism, we tested the device with dynamic assessment of cravings and two blood sampling regimes in participants at Stony Brook University School of Medicine (*Studies A, B*). To test whether the delivery method could trigger craving cycles that, in turn, were linked to drug-administration, we repeated the experiment with only dynamic cravings assessment at the Massachusetts General Hospital A.A. Martinos Center for Biomedical Imaging. This was initially done using lengthened inter-trial intervals (*Study C*), and then, using *ad libitum* self-administration (*Study D*). Finally, to confirm that the delivery method activated the reward circuit, we scanned individuals’ response to nicotine puffs using ultra-high-field/ultra-fast fMRI at the Massachusetts General Hospital A.A. Martinos Center for Biomedical Imaging (*Study E*). Characterization of subjects for each of the five studies is shown in **Table 1**.

**Table 1:**
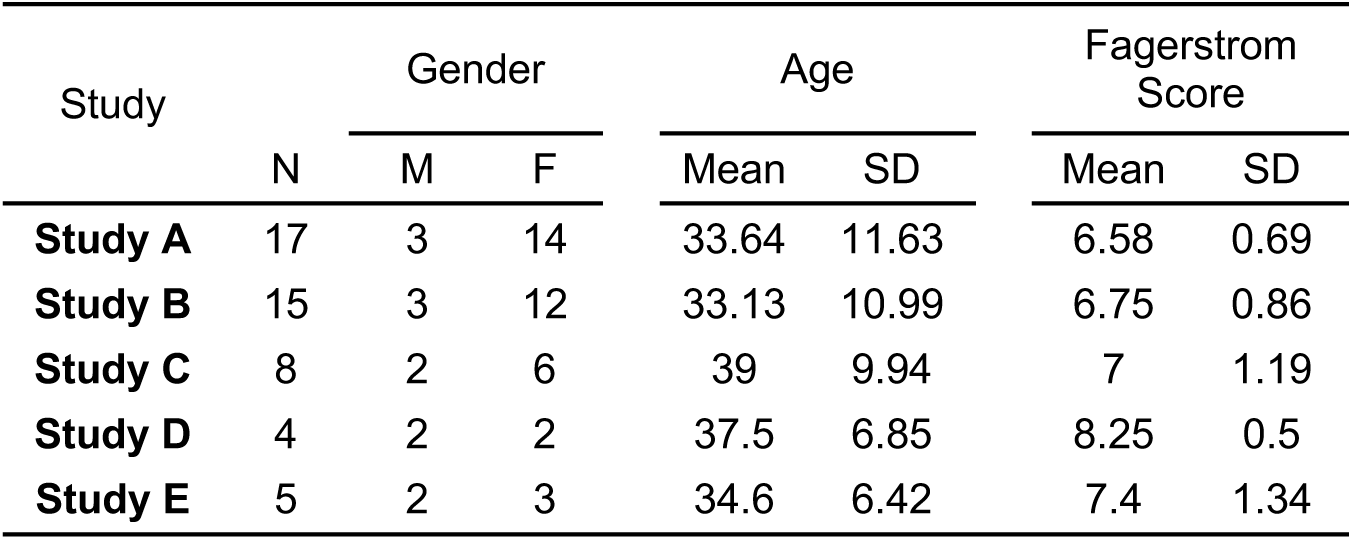
Characterization of Subjects

For all studies, we recruited subjects who were otherwise healthy daily smokers with moderate to severe nicotine dependency, and therefore who would be likely to show strong cravings following the 12-hour abstinence period preceding each session. Nicotine dependency was measured on a scale of 0-10 using the Fagerstrom Test for Nicotine Dependence (FTND) (Heatherton Todd et al. 1991). For all studies, we included only subjects with FTND scores ≥ 6, ages 21-55 with BMI 18.5-35. Abstinence prior to testing was empirically confirmed by measuring exhaled carbon monoxide levels using a Micro+ Basic Smokerlyzer (Covita), and further confirmed via blood testing. Subjects with measured baseline carbon monoxide levels > 10ppm were excluded prior to testing; any remaining subjects with baseline nicotine levels > 10ng/mL were excluded prior to analysis. Additional exclusion criteria included nasal congestion, sinusitis, use of nicotine cessation therapy medications, history of asthma, cardiovascular, or peripheral vascular disease, history of neurological disease, or the use of psychotropic medications. These criteria would influence the absorption of nicotine through the nasal mucosa, its distribution throughout and dissipation from the blood stream, and its effect on specific brain regions (Arora, Sharma, and Garg 2002). Additional exclusion criteria concerning MR safety included electrical implants, ferromagnetic implants, claustrophobia, and pregnancy. Exclusion due to pregnancy was determined using a urine pregnancy test (Detector hcg, Immunostics Inc). Studies were approved by the Institutional Review Boards of Stony Brook University and Massachusetts General Hospital. All subjects provided written informed consent.

### Task Design

For all studies, subjects were nicotine-addicted and had abstained for at least 12 hours (validation methods described above). Immediately prior to the start of the testing session, we administered 4% Lidocaine HCl (Roxane Laboratories) in the subject’s nasal cavity to alleviate the irritation from Nicotrol delivery. Lidocaine was delivered using an intranasal mucosal atomization device (Mountainside Medical Equipment Incorporated, Marcy NY). Behavioral trials used a 1mL Lidocaine dose, while MRI trials used a 3mL dose due to the increased length of the testing session. The experimental task was developed using MATLAB and Psychtoolbox-3. Behavioral task sessions ran for 60 minutes, divided into five minutes of baseline measurements, 40 minutes of the main task with puff delivery, and 15 minutes of end of study observation.

#### Pharmacokinetic studies

*Study A* provided estimation of the entire time-course, while *Study B* was designed to capture the faster-changing dynamics during uptake. For the first, we presented 20 nicotine puffs, one every two minutes, and took seven blood samples: at baseline, after every four puffs (blood sampling every eight minutes), and 15 minutes post-administration of the last puff. For the second, we maintained the same sampling at baseline and 15 minutes post-administration of the last puff, but this time sampled after every two puffs (blood sampling every four minutes).

#### Cravings studies

To test whether the delivery method could trigger craving cycles that, in turn, were linked to drug-administration, we repeated the experiment with only dynamic cravings assessment. This was initially done using lengthened inter-trial intervals (*Study C*), and then, using *ad libitum* self-administration (*Study D*). For *Study C* we used the same 10-puff protocol, but delivered puffs every four minutes, with blood sampling only at baseline to confirm abstinence. We used longer ITIs to ensure that the reward circuit had sufficient time to resolve following each puff, thereby eliciting serial craving cycles without ever achieving full satiety. For *Study D*, one puff was delivered at the beginning of the scan, and subjects were instructed to only request additional puffs when they could no longer resist their cravings. Subjects were not permitted to request a puff more frequently than once every two minutes; thus, subjects could request a maximum of 20 puffs.

#### Neuroimaging

Finally, to confirm that the delivery method activated the prefrontal-limbic-reward circuit, we measured neurobiological response to nicotine puffs using ultra-high-field/ultra-fast fMRI at the Massachusetts General Hospital A.A. Martinos Center for Biomedical Imaging (*Study E*). We used the 10-puff protocol used in *Study C*, delivering puffs every four minutes, and blood sampling only at baseline to confirm abstinence.

### Dynamic Cravings Assessment

For *Studies A-E*, throughout each 60-minute session, subjects indicated their relative cravings intensity on a dynamic Likert scale from “0” to “100,” with “0” corresponding to “no cravings” and “100” corresponding to “strongest cravings imaginable.” This scale was represented by a continuously updated line (TR=1 second), showing the subject’s cravings over the past five minutes. Subjects used a button box to adjust cravings-ratings throughout each session.

### Continuous Blood Sampling

For *Studies A-B*, arterialized venous blood samples were collected throughout each 60 minute behavioral session as a less invasive alternative to arterial sampling (Zello et al. 1990). An intravenous line was placed in the subject’s non-dominant hand, and the hand was placed in a warming box at 50°C. Each blood draw collected 4mL of blood, which was immediately transferred to a Vacuette Clotting Tube (Greiner bio-one) for serum separation. Prior to the start of the 60-minute session, the first blood draw was drawn to determine baseline nicotine and nicotine metabolite levels. Throughout the 40-minute puff delivery period, five additional blood samples were collected. In the 20-puff protocol, blood was drawn one minute after Puffs 4, 8, 12, 16, and 20. In both 10-puff protocols, blood was drawn one minute after Puffs 2, 4, 6, 8, and 10. One final blood sample was collected at the end of the 60-minute session for all behavioral protocols. During MRI trials, no heating box was used and no IV was placed, and only the first baseline blood draw was collected. Serum was separated from these samples and analyzed by ARUP Laboratories (Salt Lake City, UT) for *nicotine, cotinine*, and *trans-3’-hydroxycotinine* levels.

### Neuroimaging

#### Acquisition Parameters

Our use of ultra-high field strength, Siemens Magnetom 7 Tesla scanner with 32-channel head coil, combined with EPI acquisition parameters (TR = 802ms, TE = 30ms, 85 slices) optimized by a dynamic phantom for dynamic fidelity, were chosen to maximize single-subject level detection sensitivity of prefrontal-limbic and reward circuits(DeDora et al. 2016). Structural scans, for spatial co-registration, were acquired as multi-echo MPRAGE with 1mm isotropic voxel size at four echoes with TE1, TE2, TE3, TE4 = 1.61, 3.47, 5.33, 7.19ms, TR = 2530ms, flip angle = 7 degrees, slice gap =0.5mm and GRAPPA acceleration = 2. B0 field map images, calculated using phase differences between gradient echo images at TE = 4.60 and 5.62ms, were also acquired (TR = 723 ms, flip angle = 36°, voxel size = 1.7 × 1.7 × 1.5 mm, 89 slices) for EPI distortion correction arising due to magnetic field inhomogeneity.

#### Preprocessing

Spatial preprocessing was performed in Statistical Parametric Mapping (SPM12). Functional images were corrected for motion (rigid realignment, 6 degrees-of-freedom) and a mean functional image was calculated for each subject. These mean functional images were co-registered to high-resolution structural images followed by segmentation to generate gray matter, white matter and deformation field images. The realigned (field map corrected) functional images were normalized to Montreal Neurological Institute (MNI) EPI template with affine registration followed by a non-linear transformation (between average fMRI and EPI template). Finally, the images were smoothed with a Gaussian kernel of 8 mm at full width at half maximum. Correction for static field inhomogeneity was performed after the realignment step using a field map-based EPI unwarping tool built in FSL(Smith et al. 2004). Nicotine is known to cause vasoconstriction and alters heart rate/blood pressure as well as respiration rate (Najem et al. 2006). The vascular effects on BOLD signal can be reduced by using nuisance regressors derived from CSF and white matter(Birn 2012; Caballero-Gaudes and Reynolds 2017) that are unlikely to show neural activity induced T2* changes. We used the CompCor method (Behzadi et al. 2007), implemented using CONN toolbox (Whitfield-Gabrieli and Nieto-Castanon 2012), to account for the effect of physiological noise on the BOLD signal. CompCor regresses out the confounding effects of multiple empirically estimated noise sources calculated from variability in BOLD time-series of cerebrospinal fluid and white matter (principal component analysis). Five components of white matter, five components of the cerebrospinal fluid and six motion parameters, along with despiking and quadratic detrending, were used for denoising. The first 10 scans were removed from each dataset before analysis to permit stabilization.

#### Independent Component Analysis

Spatial Independent Component Analysis (ICA) provides a data-driven exploratory method for revealing brain activation by separating each voxel time-series into a linear combination of spatially independent source signals, enabling estimation of neural activity and confounds without a priori information. This technique is essentially assumption-free and, as such, can be more appropriate than GLM analyses in cases (as with pharmacological challenge) where the standard hemodynamic response function may not accurately model the brain’s response or is unknown(McKeown, Hansen, and Sejnowski 2003). Spatial ICA can reveal functional activity obscured from traditional GLM analysis (Xu, Potenza, and Calhoun 2013) and has proved not only to have higher specificity to activation but also higher sensitivity in analyses of ultra-high field fMRI data (Robinson et al. 2013).

The ten puff (fixed interval) protocol, delivering a puff every four minutes as in study C, was used during neuroimaging. Given the cyclic behavior of cravings and satiety established in *Study C*, and previously established regions associated with the addiction circuit(Koob and Volkow 2010), the preprocessed fMRI data were analyzed to dissociate two processes. The first is acute response to nicotine administration; the second is transition from higher to lower cravings. To analyze the response to nicotine administration, we compared the baseline measurement (before delivering the first puff) to the period exactly after each puff delivery. The fMRI data were split into two windows of 370 TRs (296.74 seconds). The first window consisted of 370 TRs acquired before administration of nicotine puffs in the baseline measurement condition (BASELINE in **Figure 4**), while the second window consisted of 370 TRs obtained by concatenating 37 TRs (29.67 seconds) following each nicotine puff (PUFF in **Figure 4**). Similarly, for the second objective to identify response to cravings, we compared the neural response during craving and satiety, and accordingly the fMRI data were split into two windows of 1000 TRs (802 seconds) each. The first window consisted of 100 TRs pre (CRAVING in **Figure 4**) nicotine administration concatenated together over each nicotine puff to form a single 1000 TRs long window, while the second window consisted of 100 TRs post (SATIETY in **Figure 4**) nicotine administration concatenated together over each nicotine puff to form the other 1000 TRs long window. The 20 TRs immediately following each puff administration were omitted from the SATIETY condition to remove effects associated with the PUFF condition.

Spatial independent component analyses (ICA) (V.D. et al. 2001) was performed on each preprocessed and windowed functional MRI dataset using Group ICA of fMRI Toolbox (GIFT v3.0b) (Rachakonda et al. 2010). Subject-specific spatial component maps (Z-scaled) and associated time courses were obtained through single subject level ICA runs. For each subject, the data was decomposed into 40 independent components. ICA was run with 10 runs of ICASSO procedure to ensure component stability (Himberg, Hyvärinen, and Esposito 2004). For reducing computational complexity, data reduction was achieved by performing subject-specific principal component analysis (PCA). ICA was then performed on this final dataset using Infomax algorithm (Lee et al. 2000). Infomax algorithm extracts independent sources from mixed sources using entropy maximization principle.

#### Analyses of Response to Nicotine Administration and Cravings

To identify the components of interest, component spatial maps obtained for each subject for both baseline measurement and nicotine puffs windows were spatially sorted, based on their spatial correlation with masks for the addiction circuit (Koob and Volkow 2016). These sorted components were then intersected with the corresponding masks to extract out regions of interest. The masks for the addiction circuit were obtained from CONN Toolbox (Whitfield-Gabrieli and Nieto-Castanon 2012) which uses FSL Harvard-Oxford Atlas (maximum likelihood cortical and subcortical atlas), except for Substantia Nigra—which was obtained from the Hammersmith Atlas (Hammers et al. 2003). Statistical maps of neural response were obtained by voxel-wise subtraction of spatially sorted masked component maps for puff and baseline measurement window for each addiction circuit region, followed by voxel-wise conversion to Z value. Finally, these statistical maps were thresholded at p<0.05 with Bonferroni correction. The same procedure above was used for calculating the response to cravings using the pre and post nicotine administration windows.

#### Statistical Analysis

*Study E*’s aim was to validate a stimulus delivery method, using previously-established regions of interest associated with nicotine addiction (Koob and Volkow 2016), rather than to discover new information about the brain. As such, we conducted single subject level analysis to show efficacy of our device for probing addiction dynamics. This required only a small number of subjects (N=5), but with the stricter-than-normal requirements that only regions showing consistent *bilateral* activation or deactivation with very conservative thresholding (p<0.05, Bonferroni Corrected) were considered positive in showing a response to nicotine administration. These requirements were made possible by the increased detection sensitivity afforded by 7T field strength(Pohmann, Speck, and Scheffler 2016), as well as acquisition parameters specifically optimized by dynamic phantom for increased SNR in our targeted regions of interest(DeDora et al. 2016).

## Results

### Our nicotine delivery method showed slightly greater delivery efficiency than that of the average cigarette

An average male of 73.5 kg (162 lbs) has an average blood volume of 5.5L. Each spray delivered 0.1mg of nicotine into the nasal cavity. The highest nicotine blood concentration after one spray was therefore 0.1mg/5.5l = 18ng/ml. Fitting the data, we found that on average each spray of 10µl delivered 0.6ng/ml into the blood stream; therefore, our delivery efficiency was 3.4%. By comparison, the average cigarette contains 12mg (range is 8– 20mg). For the same 73.5 kg (162 lbs) smoker, one cigarette results in a nicotine level in the blood between 10ng/ml and 50ng/ml, with a delivery efficiency of 0.5–2.3%.

### Our nicotine delivery method showed corresponding effects on cravings

As shown in **Figure 2**, with more rapid (ITI= 2m) puff administration, cravings decreased exponentially with each puff. With an exponential fit, A+B*exp(–t/τ): A=4.2 +/– 0.07, B=3.2 +/– 0.3, and τ=310s +/– 45s. Since the nicotine delivery was linear in time, and cravings reduced exponentially, the relationship between cravings and nicotine was also exponential. Each puff delivered 0.6ng/ml into the blood, and the decay time was 310 seconds, or 2.6 puffs. In terms of nicotine, 1.55ng/ml was the exponential decay concentration. Therefore, our observed relationship with this delivery method was: *cravings* = A+B*exp(–*nicotine concentration* / 1.55ng/ml). As shown in **Figure 3**, by doubling duration between puffs (ITI=4m), we were able to achieve oscillating transitions between cravings and satiety that, for our *ad libitum* design, corresponded with behavioral self-administration of nicotine.

**Figure 2:**
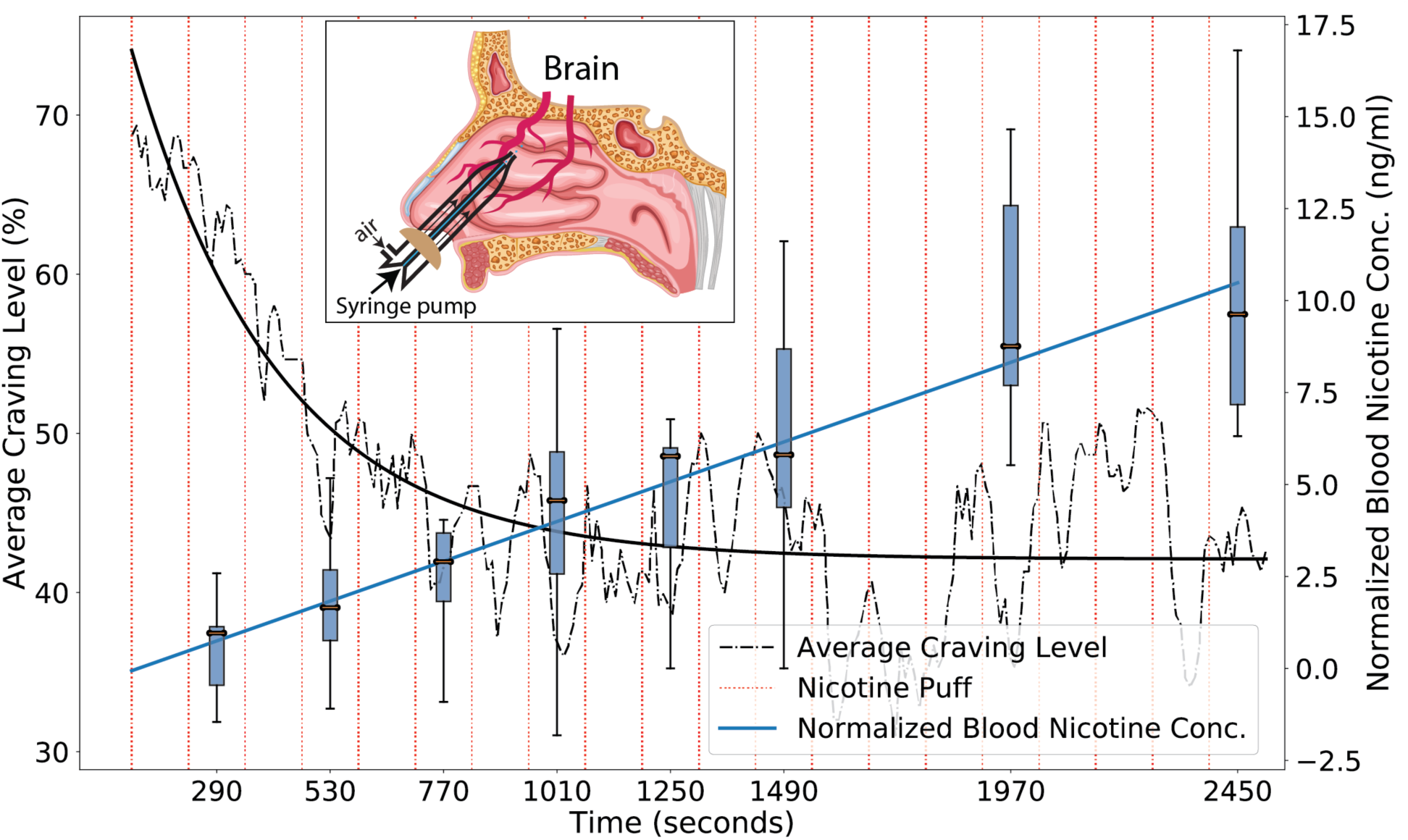
Nasal nicotine administration every 2 minutes linearly increases nicotine concentration and exponentially decreases subjective cravings. On average each spray of 10µl delivers 0.6ng/ml into the blood stream, with a delivery efficiency of 3.4% (*Studies A, B*).

**Figure 3:**
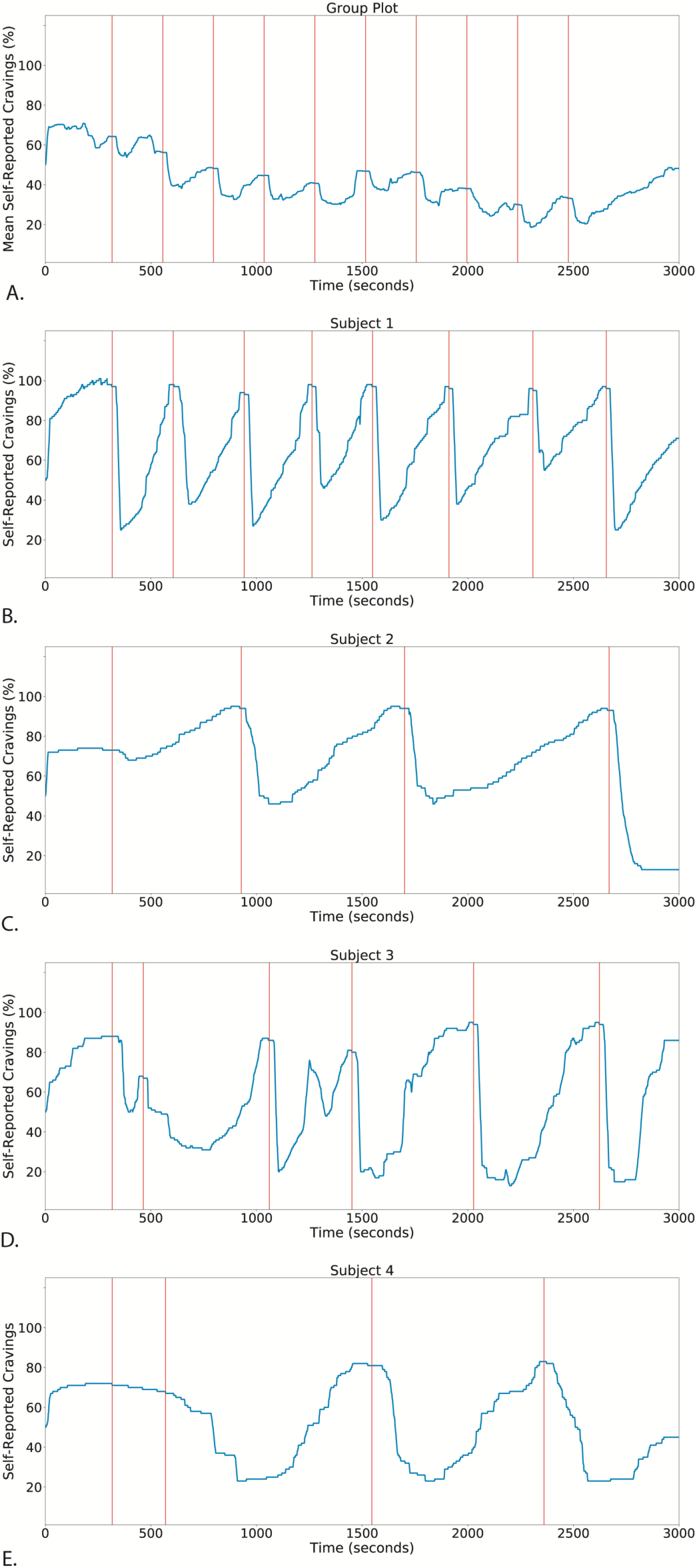
Craving Cycles. Nasal nicotine microdosing elicits craving cycles for fixed inter-trial intervals (4 minutes) **(A**, *Study C*) and predicts individual dynamics of self-administration **(B-E**, *Study D*).

**Figure 4:**
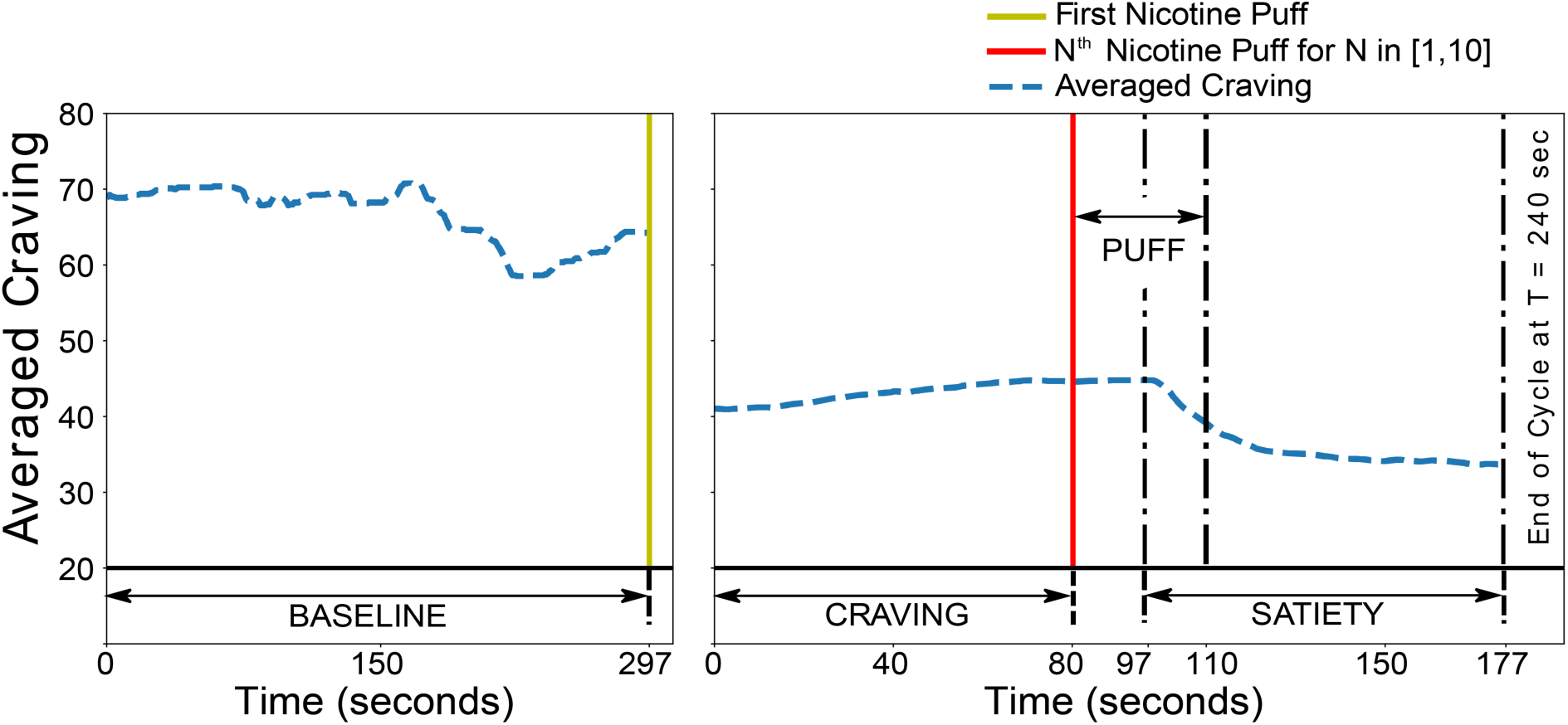
Study Design. Schematic for fMRI (*Study E*) study design components used in analysis.

### A wide network of regions shows change in activity following nicotine puffs and during transition from cravings to satiety

Our results provide significant overlap with the proposed networks described in an overview of the animal and human addiction literature (Koob and Volkow 2016), which dissociate three subcomponents of the addiction circuit, associated with reward (basal ganglia), withdrawal (amygdala), and anticipation/salience (prefrontal cortex) (note that, by design, our protocol omitted the drug-salient cues associated with the salience state that is normally included in fMRI studies of addiction). Overall, the patterns of activation can be clustered into two sets: first set containing subject 1 and subject 2, while the second containing subjects 3, 4 and 5. Specifically, nicotine administration (comparison of puffs windows with baseline) showed a decrease in activity in anterior cingulate cortex and bilateral caudate, hippocampus, putamen and ventral pallidum for subjects 1 and 2. Subjects 3, 4 and 5 show an increase in activity in most circuit regions with anterior cingulate cortex and amygdala being the common ones among all three subjects. Importantly, subject 3 and subject 5 show an increase in activity in substantia nigra following the nicotine puff **(Table 2)**. Similar patterns appear while comparing transient satiety and craving windows - subjects 1, 2 show large-scale deactivations and subjects 3, 4, 5 show activations, as reported in **Table 3**. Neural activity maps are shown in **Figure 4, 5**. A limitation of our study is its inability to generalize activations and deactivations across subjects due to its small sample size, but the study was designed as such for development of a novel nicotine delivery method rather than understanding brain activation/deactivation patterns.

**Table 2:**
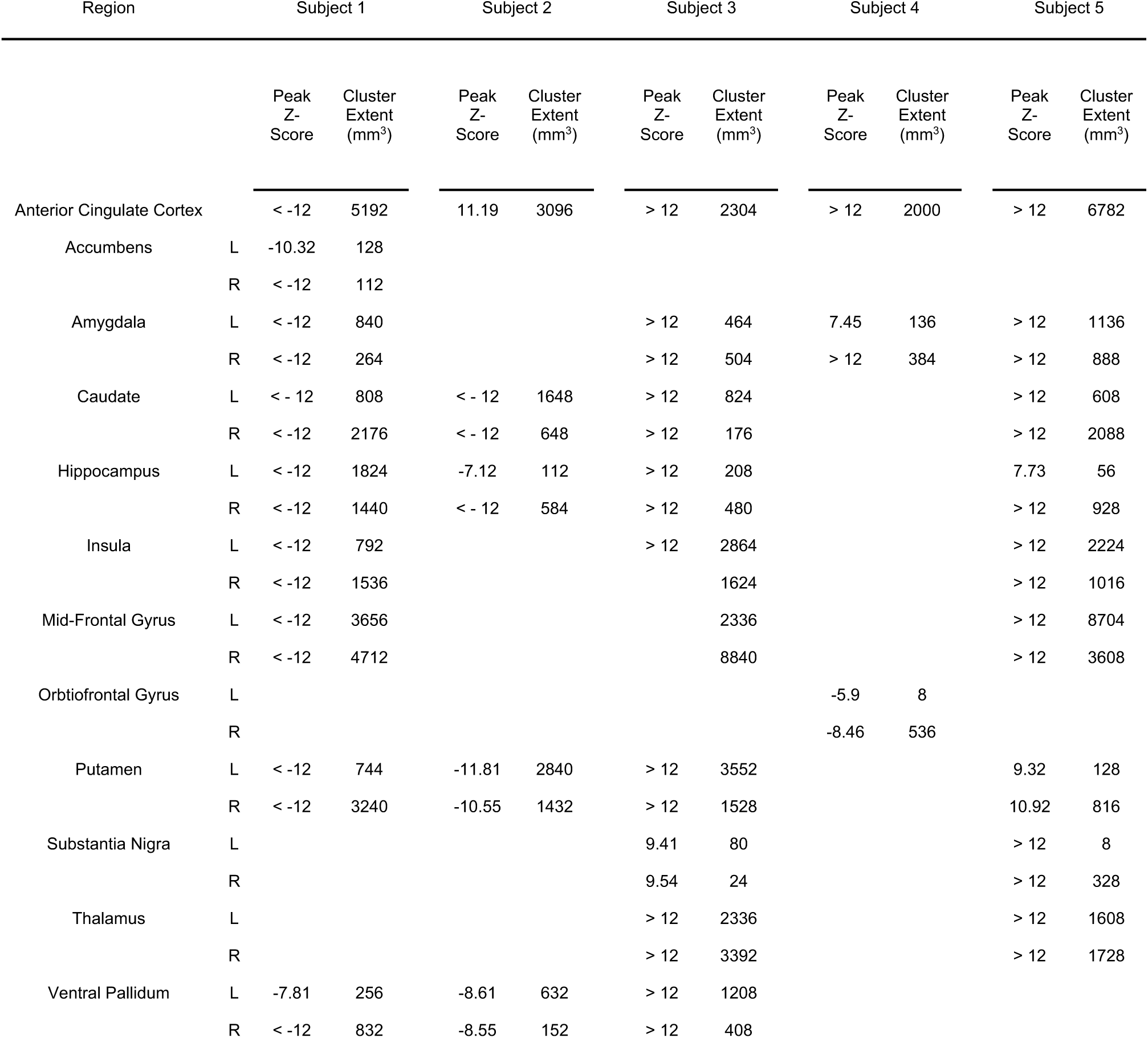
Brain Response to Nicotine Puff

**Table 3:** Brain Response to Transiently Reduced Cravings

| Region |  | Subject 1 |  | Subject 2 |  | Subject 3 |  | Subject 4 |  | Subject 5 |  |
| --- | --- | --- | --- | --- | --- | --- | --- | --- | --- | --- | --- |
|  |  | Peak Z-Score | Cluster Extent (mm<sup>3</sup>) | Peak Z-Score | Cluster Extent (mm<sup>3</sup>) | Peak Z-Score | Cluster Extent (mm<sup>3</sup>) | Peak Z-Score | Cluster Extent (mm<sup>3</sup>) | Peak Z-Score | Cluster Extent (mm<sup>3</sup>) |
| Anterior Cingulate Cortex |  | < -12 | 3704 | < -12 | 4920 | > 12 | 4176 | > 12 | 4736 | > 12 | 112 |
| Accumbens | L | -8.67 | 200 |  |  |  |  | 9.43 | 32 |  |  |
|  | R | -10.43 | 616 |  |  |  |  | 7.94 | 32 |  |  |
| Amygdala | L | -10.29 | 248 |  |  |  |  | > 12 | 584 | > 12 | 872 |
|  | R | < -12 | 128 |  |  |  |  | > 12 | 392 | > 12 | 1232 |
| Caudate | L | < -12 | 1336 | < -12 | 1344 | > 12 | 144 | 11.83 | 1296 |  |  |
|  | R | < -12 | 1000 | < -12 | 488 | 8.46 | 152 | > 12 | 536 |  |  |
| Hippocampus | L | < -12 | 896 |  |  | -7.58 | 88 | > 12 | 40 |  |  |
|  | R | -5.38 | 8 |  |  | -5.39 | 8 | 55 | 64 |  |  |
| Insula | L |  |  |  |  |  |  | > 12 | 1792 | > 12 | 3080 |
|  | R |  |  |  |  |  |  | 11.79 | 2688 | > 12 | 464 |
| Mid-Frontal Gyrus | L |  |  | > 12 | 3760 | < -12 | 7360 | > 12 | 9136 |  |  |
|  | R |  |  | > 12 | 4816 | -10.5 | 1512 | > 12 | 3728 |  |  |
| Orbitofrontal Gyrus | L |  |  | -9.64 | 968 |  |  | 9.87 | 208 | > 12 | 992 |
|  | R |  |  | < -12 | 1856 |  |  | > 12 | 1824 | > 12 | 624 |
| Putamen | L | < -12 | 2432 | -7.95 | 1344 | 6.99 | 704 | 10.1 | 632 |  |  |
|  | R | -10.37 | 1776 | < -12 | 2208 | 5.13 | 8 | 11.6 | 1312 |  |  |
| Substantia Nigra | L |  |  |  |  |  |  |  |  |  |  |
|  | R |  |  |  |  |  |  |  |  |  |  |
| Thalamus | L | < -12 | 3384 | -9.21 | 656 | > 12 | 4336 | 9.68 | 224 | -8.68 | 392 |
|  | R | < -12 | 3592 | -9.07 | 2280 | > 12 | 5560 | > 12 | 3072 | -7.55 | 896 |
| Ventral Pallidum | L |  |  | -6.31 | 120 |  |  | 6.34 | 56 |  |  |
|  | R |  |  | -8.22 | 136 |  |  | 5.57 | 24 |  |  |

**Figure 5:**
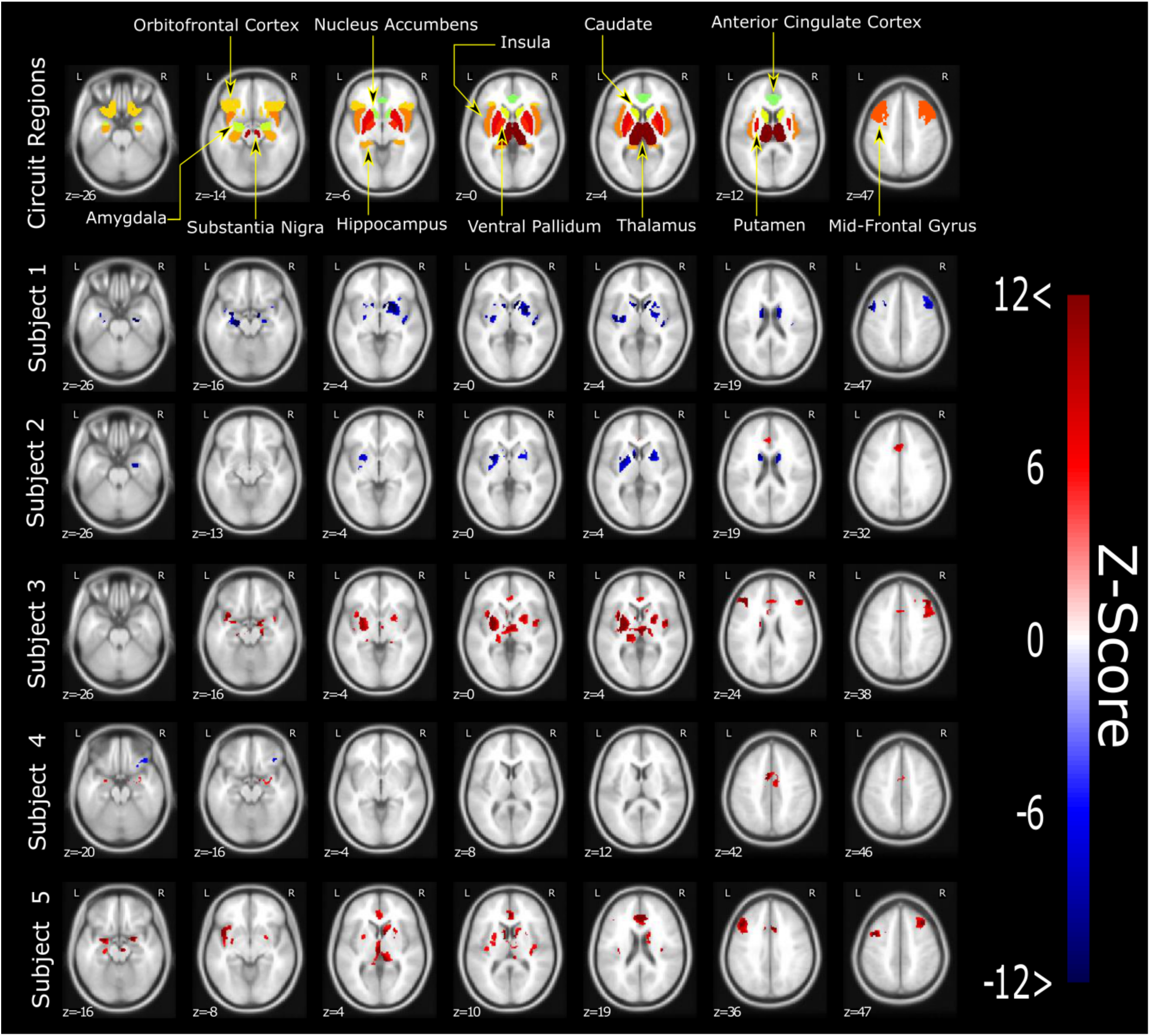
Nasal nicotine microdosing activates the addiction circuit. Composite brain activity map showing neural activity for transition between BASELINE→PUFF periods.

**Figure 6:**
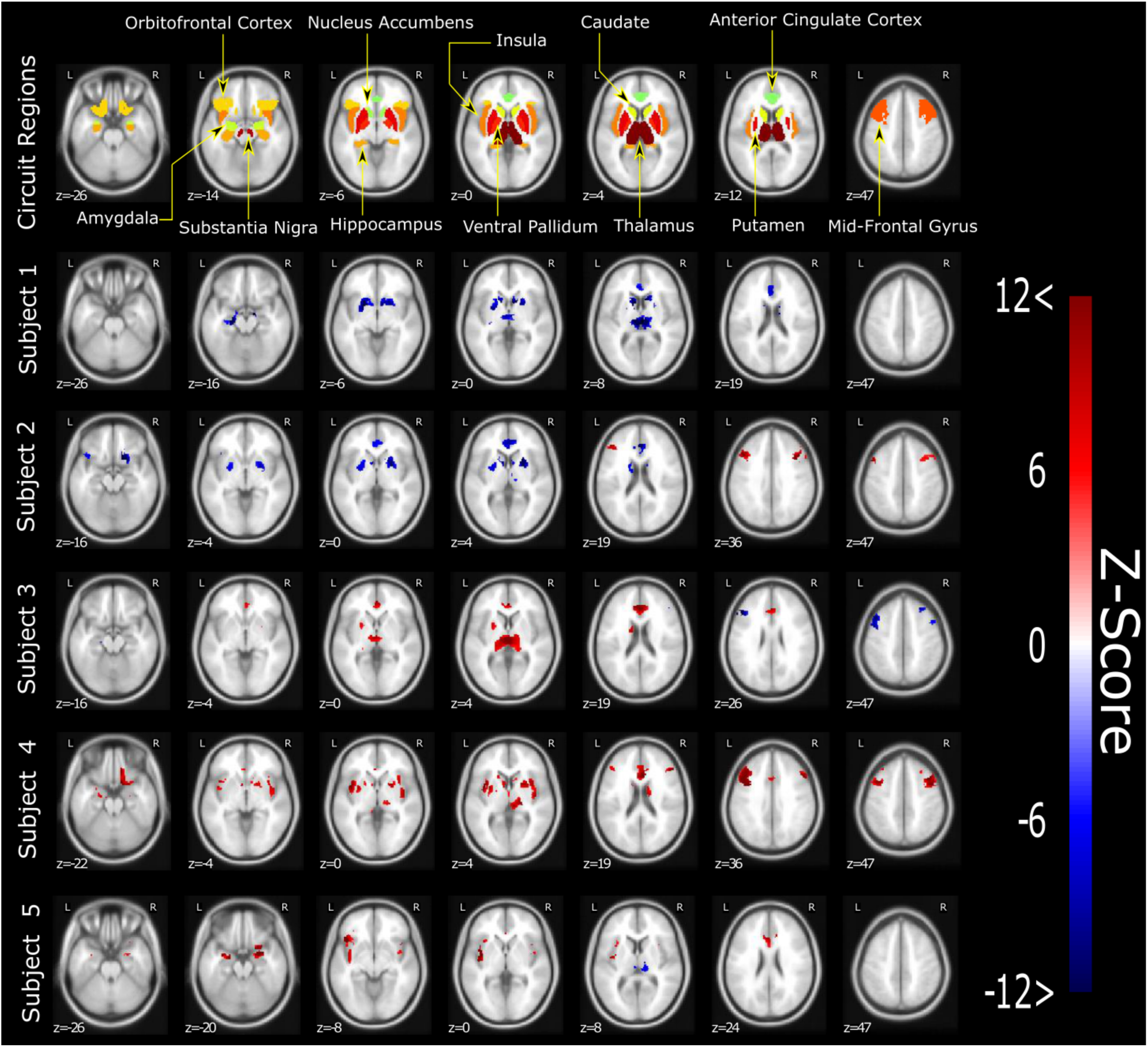
Nasal nicotine microdosing activates the addiction circuit. Composite brain activity map showing neural activity for transition between CRAVING→SATIETY periods.

Even though group-level studies are useful for discovering a generalizable set of brain regions responding to a psychostimulant, learning addiction dynamics need inference at an individual subject level (van der Stel 2015). Unfortunately, very little research has focused on single subject level analysis (Fadiga 2007), which is an absolute necessity in clinical applications. Nicotine addiction affects neural subsystems associated with decision making, emotional processing, memory, motivation, salience, and interoception – the coupled effect of which introduces variable addiction dynamics for each subject. Our study focused on an MR compatible device development which can be used for probing oscillating antagonistic sub-circuits associated with addiction. These sub-circuits modulate repeated-cycle transitions between periods of craving (affecting the nucleus accumbens and prefrontal-limbic circuit associated with aversive stimuli/emotional stress (Mujica-Parodi, Cha, and Gao 2017; LeDoux 2003; Phelps and LeDoux 2005)), reward following partial drug administration (affecting the nucleus accumbens and activating the substantia nigra subcomponents of the basal ganglia circuit (Morita et al. 2013)), and transient satiety (affecting the prefrontal-limbic circuit and the caudate and pallidum subcomponents of the basal ganglia circuit). Future efforts may be geared towards obtaining test-retest reliability of single-subject fMRI with the current delivery device and establishing associated addiction dynamics for each individual.

Our nasal drug delivery method consisted of a series of spaced micro-doses rather than a one-time dose achieving immediate or consistent saturation. This has two significant methodological advantages. First, it makes it possible to neuroimage the dynamics of self-administration, since the satiety’s transience initiates further drug-seeking behavior. Second, the cycling reveals interactions between control sub-circuits necessary for computational modeling. Importantly, the delivery device and protocol can be adapted for use with other addictive drugs (e.g., cocaine, opiates such as remifentanil with high potency and short half-life) by modulating dosage and inter-trial intervals, thereby providing a quantitative comparison of how neurobiological control dynamics, and their associated self-administrated behavior, differ across compounds. This will be particularly useful as clinical neuroimaging of addiction evolves beyond broad conceptual schemas, towards data-driven predictive models designed to rigorously quantify dysregulation and generate relapse trajectories at the single-subject level—a potential first step towards addiction-related personalized medicine.

## Funding

Research was funded by the National Institute on Drug Abuse (NIDA) Award # 3R2DA3846702S1 (LRMP).

## Conflict of Interest

The authors declare that the research was conducted in the absence of any commercial or financial relationships that could be construed as a potential conflict of interest.

## Acknowledgements

We gratefully acknowledge and thank Joanna Fowler, Lorna Role, and Edythe London for valuable scientific discussion in developing the study and Mohanlall Narine, and Sabeen Rizwan, for their assistance in collecting data.

## Notes

### Competing Interest Statement

The authors have declared no competing interest.

